# Phosphoproteomics highlights complex resource management upon inflammatory stimulation of fibroblasts

**DOI:** 10.1101/2024.10.11.617591

**Authors:** Patricia Janker-Bortel, Ana Martinez-Val, Gerhard Hagn, Lukas Skos, Jesper V. Olsen, Christopher Gerner

## Abstract

The stimulation of cells by inflammatory mediators gives rise to intricate signaling cascades, inducing specific biological functions. The role of kinase activation for the establishment of specific inflammatory functions has only scarcely been established. A time-course analysis of human fetal fibroblasts stimulated with Interleukin-1β (IL-1β) and/or Dexamethasone (Dex) was conducted using mass spectrometry-based proteomics and phosphoproteomics in conjunction with lysolipid and oxylipin profiling. The IL-1β induced proteome alterations indicated metabolic, transcriptional, and translational activation, including inflammatory marker proteins such as NOS1, THBS1, and STING1. The induction of mitochondrial proteins and the formation of numerous lysolipids indicated an increase in beta-oxidation. In addition to the NF-κB and STAT signaling pathways, which are characteristic of inflammatory activation, the MAP and AKT kinase signaling pathways were found to be strongly induced. Six hours after treatment, the observed signaling events exhibited a notable decline, nearly returning to their initial states after 24 hours. It is noteworthy that nearly all of these signaling activities were also observed in cells treated with Dex alone. Additionally, the proteome exhibited transient alterations, which included proteins otherwise characteristic of an inflammatory response, such as MMP3 and NFKB2. Activation of the kinase PIKFYVE was apparently specific for dexamethasone but constituted a minority of the total phosphorylation events. Only after 24 hours was the induction of proteins characteristic of glucocorticoids, such as TSC22D3 and MAOA, observed. The analysis of the effects of dexamethasone on the background of established inflammatory signaling verified the known inhibitory functions and resulted in the expression of anti-inflammatory proteins and oxylipins, but hardly affected the signaling events involving MAP and AKT kinases. In conclusion, this data demonstrates that a majority of inflammation-associated signaling events in fibroblasts needs to be attributed to resource and stress management rather than the establishment of specific inflammatory effector functions.

## INTRODUCTION

Inflammation is a fundamental biological process playing a crucial role in a broad variety of physiological processes such as host defense, resolution of infections and tissue repair.^1–3^ However, persistent and dysregulated inflammatory response can lead to chronic inflammation, which is increasingly recognized as a key driver in the pathogenesis of numerous diseases^4–6^, including cancer^7^, fibrosis^8^, atherosclerosis^9^, inflammatory bowel diseases (IBDs)^10^, obesity^11^ and rheumatoid arthritis (RA)^12^.

Fibroblasts, traditionally viewed as homogeneous stromal cells, are nowadays understood to include highly diverse cell subtypes^13–15^ and to play an important role in wound healing, extracellular matrix remodeling and immune response.^16,17^ Being involved in acute and chronic inflammation, they are a key cellular driver of chronic tissue inflammation in diseases such as IBDs and RA *via* their release of inflammatory cytokines and chemokines.^18,19^ This emerging evidence underscores the importance of understanding their response to inflammatory stimuli, their inflammatory status and the signaling pathgways and networks underlying their functional response.^20^

While established therapeutic approaches might target fibroblast-derived cytokines, there are currently no FDA-approved therapies directly targeting fibroblasts in inflammatory diseases.^17^ Although significant process has been made in characterizing many key signaling pathways activated in response to inflammatory stimuli, ongoing research continues to reveal so far unnoticed regulatory mechanisms and nuances that contribute to the complexity of these processes.^21–23^

The dynamic and often transient nature of these signaling events presents a significant challenge. Traditional approaches, such as the use of antibodies to investigate signaling pathways, have been widely used and provided valuable insights, but are often limited by their rather targeted nature and the inability to capture the global landscape of phosphorylation events.^24–26^ In contrast, mass spectrometry-based phosphoproteomics offers a powerful and comprehensive approach to map kinase signaling cascades in an unbiased manner^26,27^, eventually providing a broader and more detailed perspective on the cellular processes involved in inflammation.

Recent advances in mass spectrometric (MS) instrumentation and phosphoproteomics sample preparation methods have enabled the quantification of thousands of phosphorylation sites in a single experiment^28–31^, revealing new regulatory mechanisms and providing insights into the spatial and temporal dynamics of signaling networks^32–35^. The integration of proteomics and phosphoproteomics with other omics techniques, such as eicosadomics, which profiles bioactive lipid mediators like oxylipins and lysolipids, could further enhance our understanding of the molecular players involved in inflammation^10,16,36^. Such a combined multi-omics approach might not only allow for the identification of so far unnoticed signaling events but could also shed light on the metabolic processes that support and modulate the inflammatory response.

In this study, we employed a time-course analysis using MS-based proteomics, phosphoproteomics and eicosadomics to investigate the signaling and metabolic responses of human embryonic skin fibroblasts to IL-1β stimulation. Furthermore, we examined the effects of dexamethasone, a synthetic glucocorticoid widely used for its anti-inflammatory properties.^37^ Dexamethasone exerts its effects primarily through the glucocorticoid receptor, modulating a range of cellular processes^38,39^ and down-regulating pro-inflammatory cytokines^16,40^. This study provides temporally resolved insights into the cellular strategies employed by fibroblasts to manage inflammatory and anti-inflammatory stimuli, as well as the resolving of inflammation.

### EXPERIMENTAL PROCEDURES

### Experimental Design and Statistical Rationale

All experiments in this manuscript were performed as biological replicates (n=4).

#### Cell Culture

##### Culture Conditions

The skin fibroblast cell line Detroit 551 was purchased from ATCC (Manassas, VA, USA). Detroit 551 cells were cultured in Eagle’s minimum essential medium (EMEM; Sigma–Aldrich, USA) supplemented with 10% fetal calf serum (FCS) and 1% penicillin/streptomycin (Gibco, USA). The cells were grown in a humidified atmosphere with 5% CO_2_ at 37°C.

##### Treatment

Detroit 551 cells were cultured in T75 flasks for adherent cell culture (Sarstedt GmbH, Nümbrecht, Germany) until approximately 75% confluence was reached. For IL-1β and IL-1β + Dexamethasone (Dex) treated samples, the medium was exchanged with medium containing IL-1β at 10 ng/mL. For samples of the Control (Con) and Dex group, the medium was exchanged with a fresh aliquot. After 1h of treatment with IL-1β, 100µL of 0.1% [vol/vol] dimethyl sulfoxide (DMSO) were added to the medium (10 mL per flask) of the Con and the IL-1β group as solvent control. 100 µL of 0.01 µg/µL Dexamethasone dissolved in DMSO were spiked into the medium of the samples of the Dex and IL-1β + Dex group to achieve a final concentration of 100 ng/mL. Treated cells were harvested after 2 h, 6 h and 24 h total incubation time starting from the first medium exchange. Cells of the control group were harvested after 24 h. All conditions were prepared in quadruplicates.

##### Sample Work-up

All steps for sample collection were performed on ice. The supernatant of each flask (10 mL) was transferred into a 15 mL Falcon® tube and centrifuged to remove cellular debris (1233 g, 5 min, 4 °C). 3 mL of the resulting supernatant were added to 12 mL ice-cold ethanol containing 5 µL of an internal eicosanoid standard mixture (Cayman Chemical, Tallinn, Estonia) (**Supplementary Table S1**). The mixture was inverted once and stored at – 20°C overnight before further processing according to the oxylipin and fatty acid workflow. The Detroit 551 cells were washed twice with 3 mL of cold Tris-buffered saline (TBS; 25 mM Tris·HCl, 150 mM NaCl, pH 7.4). The buffer was removed completely and 200 µL of lysis buffer (5% [wt/vol] SDS, 5 mM Tris(2-carboxyethyl)phosphine hydrochloride (TCEP), 10 mM chloroacetamide (CAA), 100 mM Tris pH 8.5) were added. Cells were scraped, the whole cell lysates (WCLs) were transferred to Covaris® tubes and the lysates were immediately heat-treated for 10 min at 95°C. WCLs were homogenized using the S220 Focused-ultrasonicator (Covaris, LLC., Woburn, MA, USA), employing a lysis method with 140 W peak incident power (PIP), 10% duty factor and 200 cycles per burst (cpb) for a total runtime of 120 s at 15°C. Homogenized cell lysates were transferred to Protein LoBind® Tubes and stored at –80°C until further processing *via* the proteomics and phosphoproteomics workflow.

#### Proteomics & Phosphoproteomics

##### Sample Preparation

Samples were thawed at 37°C for 15 min, cooled to room temperature and centrifuged (10 000 g, 5 min). Protein concentration was determined utilizing the Pierce™ BCA Protein Assay Kit (Thermo Scientific™, Waltham, MA, USA) according to the manufacturer’s instructions for the 96-well plate setup.

##### Automatized Protein Aggregation Capture (PAC)-Based Protein Digestion

Protein digestion was performed using a modified version^30^ of the Protein Aggregation Capture (PAC) based digestion protocol^41^ on a KingFisher™ Flex System^42^ (Thermo Scientific) utilizing MagReSyn® Hydroxyl beads (ReSyn® Biosciences™). For the washing steps, KingFisher™ 96 deep-well plates were prepared containing 1 mL 95% Acetonitrile (ACN) or 70% Ethanol (EtOH). 300 µL of digestion solution (50 mM ammonium bicarbonate (ABC)-buffer) containing Trypsin (Sigma Aldrich) and Lys-C (Wako) at an enzyme-to-substrate ratio of 1:250 and 1:500, respectively, were prepared for each sample and transferred to KingFisher™ plates. Sample volumes corresponding to 500 µg of protein were mixed with 100 mM Tris-buffer to achieve a total volume of 300 µL. The diluted samples were transferred to KingFisher™ plates and ACN was added to achieve a final volume percentage of 70%. The storage solution was removed from the Hydroxyl beads and replaced with 70% ACN before beads were added to the samples at a protein-to-beads ratio of 1:2. Protein aggregation was performed in two steps of medium mixing for 1 min, followed by a 10 min pause each. Slow washing was performed for 2.5 min without releasing the beads from the magnet, followed by digestion within 100 cycles of 45 s agitation with 6 min pauses overnight at 37°C. Protease activity was quenched by acidification with trifluoroacetic acid (TFA).

##### Sep-Pak Desalting and Determination of Peptide Concentration

Desalting of digested samples was performed on Sep-Pak C18 Cartridges (C18 Classic Cartridge, Waters), followed by determination of the peptide concentration as described previously^30^. Aliquots of the peptide solutions were taken for proteome analysis, diluted with 0.1% FA and stored at –20°C until further processing.

##### Phosphopeptide Enrichment

Phosphopeptide enrichment was performed on a KingFisher™ Flex System employing MagReSyn® Zr-IMAC HP beads (ReSyn® Biosciences™). Sample volumes containing 137 µg of peptide were mixed with 200 µL loading buffer (80% ACN, 5% TFA, 0.1 M glycolic acid) and transferred to a KingFisher™ plate. Further plates containing 500 µL of loading buffer, 500 µL of washing buffer 2 (80% ACN, 1% TFA), or 500 μL of washing buffer 3 (10% ACN, 0.2% TFA) per well were prepared. 14 µL of beads (20 mg/mL) were added to 500 µL of 100% ACN in a plate for each sample. 200 µL elution buffer (1% NH_4_OH) were prepared per sample. Beads were washed with loading buffer for 5 min at 1000 rpm, mixed with the samples for 20 min at medium speed and washed in loading buffer, washing buffer 2 and washing buffer 3 for 2 min each at fast speed. Phosphopeptides were eluted with elution buffer by mixing for 10 min at fast speed. Eluates were acidified with 40 µL of 10% TFA to achieve a pH <2.

##### Sample Preparation for LC-MS/MS Analysis

Acidified phosphopeptide eluates were transferred to Multi-Screen_HTS_-HV 96-well filtration plates (0.45 µm, Millipore), stacked on 96-well plates and centrifuged (500 g, 1 min) to remove in-suspension particles. Desalted peptide samples for proteome analysis were thawed and spun down. Proteome and phosphoproteome samples were loaded onto Evotip Pure™ (Evosep) according to the manufacturer’s instructions.

##### LC-MS/MS Analysis

Proteome and phosphoproteome samples were separated on an EvoSep One LC system (Evosep) using the pre-programmed gradient for 30 samples per day (SPD) employing a EV1137 column (C18, 15 cm × 150 µm, 1.6 µm, Evosep). The column temperature was maintained at 50 °C using a butterfly heater (PST-ES-BPH-20, Phoenix S&T) and interfaced online using an EASY-Spray™ source with the Orbitrap Exploris 480 MS (Thermo Fisher Scientific, Bremen, Germany).

The spray voltage was set to 1.8 kV, the funnel RF to 40 and the heated capillary temperature to 275°C. The full MS resolution at m/z 200 was set to 120,000, the full MS AGC target was 300% and the maximum injection time (IT) 45 ms. The MS2 resolution was set to 15,000, the IT to 22 ms, the normalized collision energy (NCE) to 27% and the AGC target value to 1000%. Data was acquired *via* data -independent acquisition (DIA). For phosphoproteome samples, the MS2 mass range was 472 to 1143 m/z. For proteome samples, the MS2 mass range was 361 to 1033 m/z. The mass ranges were scanned within 49 windows of 13.7 m/z width with an overlap of 1 Da within a total cycle time of 2 s. The total runtime per sample was 45 min.

##### Data Processing

Raw files of proteome and phosphoproteome samples were searched using Spectronaut™ (version 17.1) using the directDIA+™ (Deep) search strategy against the Homo sapiens proteome retrieved from the UniProt Database^43^ (version 2022, 20,958 entries) supplemented with a database of common contaminants (246 entries). Cysteine carbamidomethylation was set as fixed modification. Variable modifications included N-terminal protein acetylation and oxidation of methionine for proteome samples, as well as phosphorylation of serine, threonine and tyrosine for phosphoproteome samples. The MS2 de-multiplexing was set to automatic. The digest type was set to specific with Trypsin/P as enzyme and a maximum of two missed cleavages per peptide. A maximum of five variable modifications per peptide were allowed. Method evaluation mode was turned off. Only for phosphoproteome samples, the PTM localization was enabled, but the localization probability threshold was set to 0. The mass tolerance strategy settings were default, both for MS1 and MS2, and the peptide spectrum match (PSM), peptide and protein group FDR were set to 0.01. The precursor q-value and the posterior error probability (PEP) value were set to 0.01 and 0.2, respectively. The protein level q-value cutoffs were set to 0.01 experiment wide and 0.05 run wide. The mutated strategy was used for decoy generation. Quantification was performed area-based at MS2 level, with precursor filtering set to “Identified (Q value)”. Cross-run normalization was turned off, no imputation was performed at this step and the results were filtered for the best 3-6 fragments per peptide. All remaining parameters were set to default.

For proteome samples, a standard format report was exported from Spectronaut. For phosphoproteome samples, a PTM report in long-format was exported from Spectronaut and imported into Perseus^44^ (version 1.6.5.0). Using the “peptide-collapse” plugin^45^ (version 1.4.2), the data was collapsed into phosphosites at the “Target PTM site-level” with a localization probability cutoff of 0.75.

Further data processing was performed in RStudio^46^ (version 2024.4.0.735) employing R^47^ (version 4.3.3). LFQ intensity values were log_2_ transformed and filtered for at least 3 out of 4 valid values in at least one experimental group. Data normalization was performed using the “normalizeCyclicLoess” function from the package limma^48^ (version 3.58.1). For proteomic data, missing values were imputed with the “impute.slsa” and the “impute.pa” function from the package “imp4p”^49^ (version 1.2) depending on whether they were missing at random, or missing at not random. For phosphoproteomic data, missing values were imputed using the “scImpute” and the “tImpute” function from the package “PhosR”^50^ (version 1.12.0) and data was centered across their median. Differentially expressed proteins and phosphosites were determined with the “lmFit” and “eBayes” function from limma^48^ (version 3.58.1). Adjusted p-values were calculated employing the default Benjamini-Hochberg method. Proteins with an adjusted p-value ≤0.05 and a log2 fold-change ≤–0.3 or ≥0.3 as well as phosphosites with an adjusted p-value ≤0.05 were considered as significantly regulated.

##### Data visualization

If not stated otherwise, data visualization was performed using the R package “ggplot2”^51^ (version 3.5.0). Heatmaps and UpSet plots were generated using the R package “ComplexHeatmap”^52^ (version 2.18.0).

##### CausalPath analysis

Proteomics and phosphoproteomics data were integrated into causal networks utilizing CausalPath^53^. The LFQ-intensity values pre-processed as described above were used as input and standard parameter settings were applied (value transformation = significant-change-of-mean, FDR threshold = 0.1 for protein and phosphoprotein). The respective networks were visualized in Newt^54^.

##### Kinase-substrate enrichment analysis (KSEA)

A kinase-substrate enrichment analysis was performed using the phosphosites pre-processed as described above as input for the KSEA App^55^, employing PhosphoSitePlus^56^ and NetworKIN^57^.

#### Oxylipin & fatty acid analysis

##### Sample Preparation

Oxylipin and fatty acid analysis was performed as described previously^58^. In short, precipitated supernatants in ethanol were centrifuged (4536 g, 30min, 4°C) and the new supernatants were transferred to new tubes. The EtOH volume was reduced *via* vacuum centrifugation at 37°C until the original sample volume was restored. The samples were loaded onto pre-conditioned StrataX solid-phase extraction (SPE) columns (30 mg/ 1 mL, Phenomenex, Torrance, CA, USA) and subsequently washed with ice-cold MS-grade water. Analytes were eluted with ice-cold MeOH containing 2% formic acid (FA). The eluates were dried under a gentle nitrogen stream at room temperature. Dried samples were reconstituted in 150 µL reconstitution buffer (64.8% H_2_O, 31.5% ACN, 3.5% MeOH, 0.2% FA).

##### LC-MS/MS analysis

Analytes were separated using a reversed-phase Kinetex® C18 column (2.6 μm XB-C18, 100 Å, LC column 150 × 2.1 mm, Phenomenex, Torrance, CA, USA) on a Vanquish™ UHPLC system (Thermo Fisher Scientific™, Vienna, Austria) at a flow rate of 200 µL/min and at 40°C. All samples were measured in technical duplicates with an injection volume of 20 µL. A flow gradient starting at 35% mobile phase B (89.8% ACN, 10% MeOH, 0.2% FA) in mobile phase A (99.8% H_2_O, 0.2% FA) was applied and increased to 90% mobile phase B over 10 min. Mobile phase B was further increased to 99% within 0.5 min and kept for 5 min before decreasing it to 35% within 0.5 min and holding it for 4 min. The total runtime was 20 min. Sample analysis was performed on a QExactive™ HF hybrid quadrupole mass spectrometer (Thermo Fisher Scientific™) equipped with a HESI source. The MS was operated in negative ionization mode with a spray voltage of 3.5 kV and a capillary temperature of 253°C. Auxiliary gas and sheath gas were set to 10 and 46 arbitrary units, respectively. MS1 spectra were acquired in the range of 250–700 m/z with a resolution of 60,000 at 200 m/z. For MS2 spectra, a data-dependent acquisition (DDA) approach with a Top2 method was employed, using a resolution of 15,000 at 200 m/z and a NCE of 24. Additionally, an inclusion list with 33 precursor masses specific for eicosanoid and their precursor fatty acids was utilized. (**Supplementary Table S2**).

##### Data Processing

Raw files were manually curated using the Xcalibur™ Qual Browser software (version 4.1.31.9, Thermo Fisher Scientific™, Bremen, Germany), employing reference spectra from the LIPIDMAPS depository library^59^ (version 07/2022). Criteria for analyte identification included the exact analyte mass, retention time and MS2 fragmentation pattern. The TraceFinder software (version 4.1, Thermo Fisher Scientific™, Bremen, Germany) was used for relative quantification, allowing a maximum mass deviation of 5ppm. Analyte peak areas were further processed in RStudio^46^ (version 2024.4.0.735) employing R^47^ (version 4.2.0). After log_2_-transformation, the mean peak areas of the internal standards were subtracted from the analyte peak areas for normalization. Peak areas were increased by addition of (x+20) to obtain a data distribution similar to those in label-free quantification (LFQ) based proteomics. Missing value imputation was performed using the minProb function of the imputeLCMD package^60^ (version 2.1). Differentially regulated analytes were determined by using an unpaired T-test (p-value cutoff 0.05) and applying the Benjamini-Hochberg procedure as multiple testing correction.

## RESULTS AND DISCUSSION

### Experimental design to investigate signaling events involved in inflammatory functions

Inflammatory responses were triggered in human fibroblast Detroit-551 cells by supplementing the growth medium with 10 ng/mL Interleukin-1β (IL-1β). Cells were harvested at 2, 6 and 24 hours post-stimulation to generate a temporally resolved profile of the inflammatory response. To assess cellular responses to anti-inflammatory intervention, cells were pre-treated with 10 ng/mL IL-1β for 1 hour, followed by the addition of Dexamethasone (Dex) at a final concentration of 100 ng/mL. This combined treatment included IL-1β stimulation for 1 hour, followed by Dex treatment for 5, 6 and 23 hours. Additionally, to examine the effects of glucocorticoid receptor activation in the absence of prior inflammatory activation, treatment with 100ng/mL Dex alone was conducted in a matched time scheme for 1, 5 and 23 hours. For all conditions, biological quadruplicates were employed and cellular responses were monitored applying a multiomics workflow, including oxylipin and fatty acid analysis as well as proteomic and phosphoproteomic profiling as depicted in **Figure 1**.

**Figure 1.**
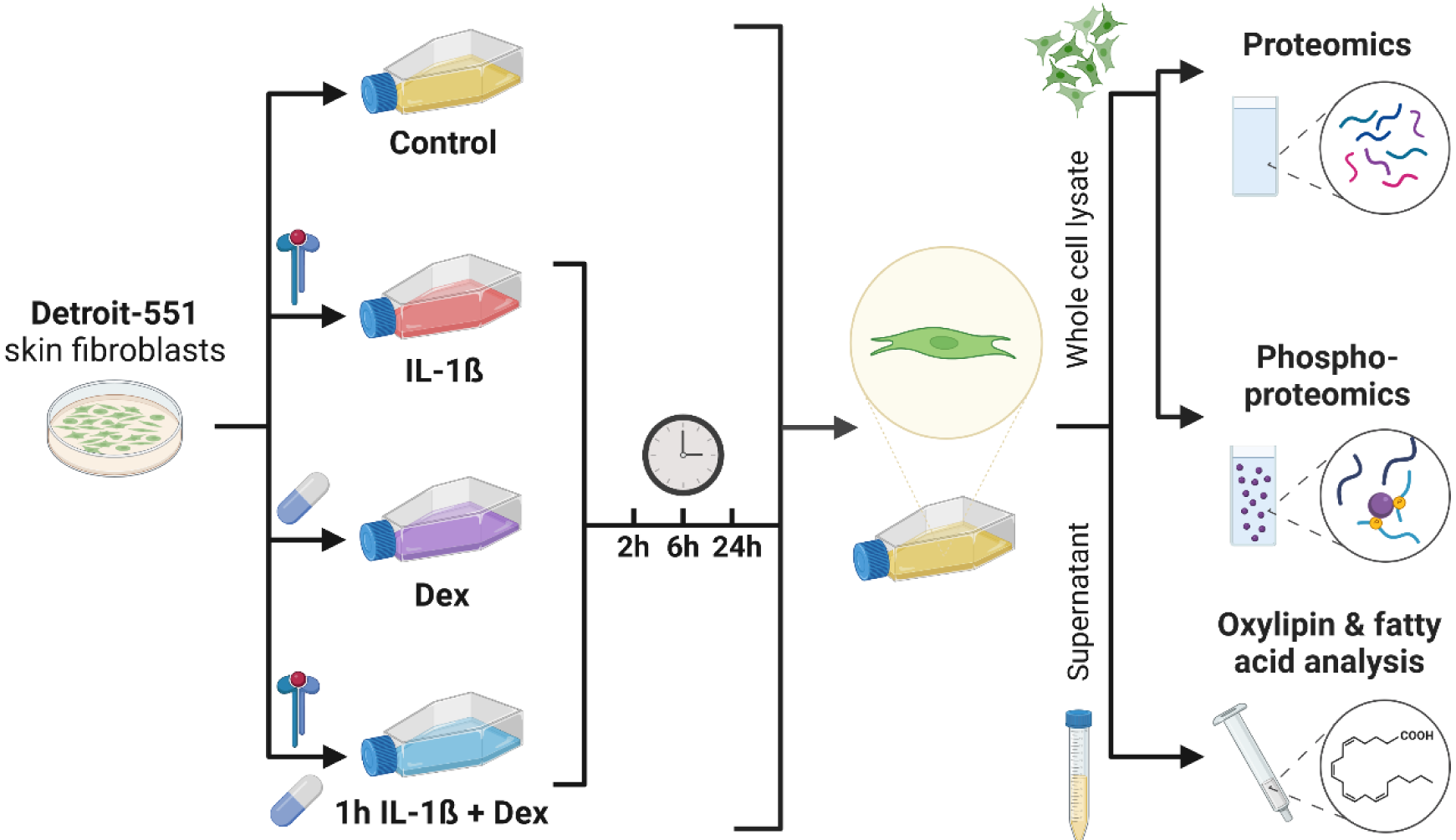
Experimental design to assess the temporally resolved proteomic, phosphoproteomic and eicosadomic profiles upon Interleukin-1β and Dexamethasone treatment on skin fibroblasts. Detroit-551 skin fibroblast cells were cultured and treated either with solvent control (“Control”), Interleukin-1β at 10 ng/mL (“IL-1β”), Dexamethasone at 100 ng/mL (“Dex”) or Interleukin-1β and Dexamethasone (“1h IL-1β + Dex”) in matched concentrations. For the combined treatment, samples were treated with IL-1β for 1 h before addition of Dexamethasone. Cells were harvested after 2, 6 and 24 h of total incubation time. All samples were subjected to a multi-omics workflow including the analysis of oxylipins and fatty acids from supernatants, and the analysis of proteins and phosphoproteins from the whole cell lysates.

### Interleukin-1β triggers a transient inflammatory signaling cascade in human skin fibroblasts

Global proteome profiling of whole cell lysates revealed numerous proteome alterations 2, 6 and 24 hours following IL-1β treatment (**Figure 2A**). Principle component analysis (PCA) of the global proteome demonstrated the greatest divergence from untreated cells already 2 hours after activation, and the smallest distance after 24 hours, suggesting transient effects of inflammatory activation, predominantly represented by PC1 (**Figure 2C**). Among the dynamically regulated proteins were inflammatory markers including MMP3^61,62^, VCAM1^63,64^, THBS1^65,66^ and NOS1^67,68^, alongside GRN^69,70^, CCN1^71,72^, STING1^73,74^ and the kinase RIPK2^75,76^, described to modulate immune responses (**Figure 2A**). Transcriptional activation was indicated by the upregulation of NF-kappa-B (p100 NFKB2)^77^ and immediate-early genes such as the transcription complex AP-1 (JUNB and FOSB)^78^ and the transcriptional regulator EGR1^79^. Increased glucose consumption was indicated by upregulation of the glucose transporter SLC2A10. Further, enhanced cellular sensitivity to potentially synergistic signals was suggested by the upregulation of the prostaglandin E2 receptor PTGER2, the nuclear receptor NR4A1 and ryanodine receptor RYR1. Six hours after inflammatory stimulation, the transient upregulation of COX2 (PTGS2), an important promoter of pro-inflammatory lipids^80^, was observed, along with increased expression of the autophagy promoters FOXO3 and RB1CC1^81,82^. Interestingly, the negative feedback regulator of innate immune responses, TRAFD1, was also induced, potentially contributing to the early attenuation of inflammatory effector protein expression (**Figure 2A).** By 24 hours post-treatment, most IL-1β-induced proteomic alterations had subsided, indicating a rapid decline of inflammatory cell activities.

**Figure 2:**
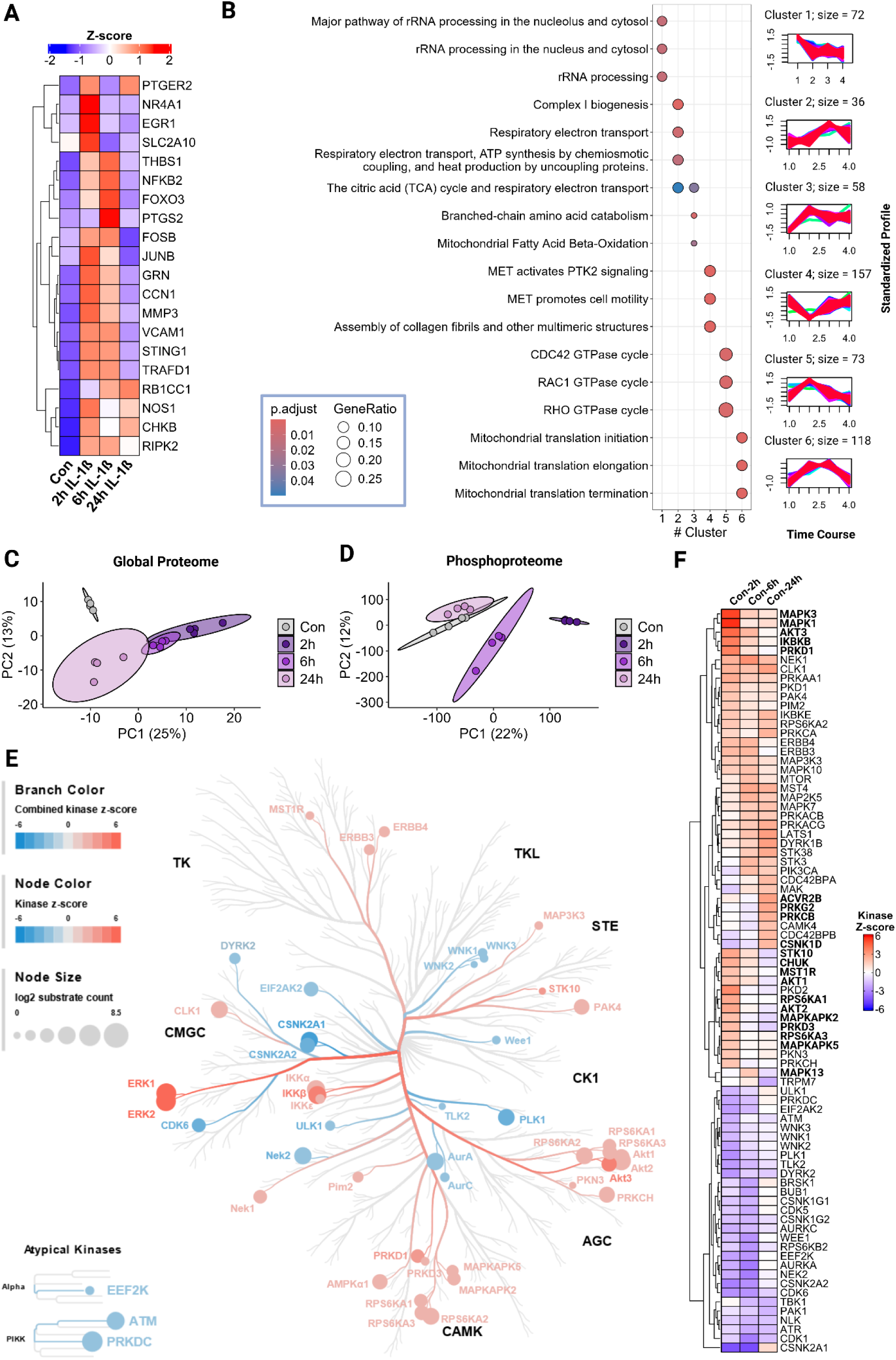
Time-resolved effects of IL-1β on the Global Proteome and Phosphoproteome and inferred kinase activities. (A) Heatmap of z-score transformed LFQ intensities of proteins described to be involved in inflammatory processes. Values for control samples and IL-1β treated samples (2, 6 and 24 h) are displayed from left to right. (B) *Right panel* – Clusters of proteins exhibiting similar temporal LFQ intensity profiles upon stimulation with IL-1β. Only proteins which were significantly regulated at at least one time point compared to the controls were subjected to the cluster analysis. *Left panel –* The dotplot illustrates the results of a Reactome pathway over-representation analysis performed with proteins contained in the respective clusters. The top 3 most significantly enriched pathways per cluster are shown. The dot color indicates the adjusted p-value of the respective enriched pathway and the dot size represents the ratio of pathway specific proteins in comparison to the total protein number contained in the respective cluster. (C) A principal component analysis (PCA) of proteome profiles of control and IL-1β stimulated samples is shown. Ellipses represent 95% confidence intervals (CI). (D) A principal component analysis (PCA) of phosphoproteome profiles of control and IL-1β stimulated samples is shown. Ellipses represent 95% confidence intervals (CI). (E) The kinome tree depicts kinases significantly enriched for substrate phosphorylations determined by performing a kinase substrate enrichment analysis (KSEA) on class I phosphosites significantly regulated upon treatment with IL-1β for 2 h. Kinase z-scores indicate the statistical significance and direction of kinase activities inferred from the KSEA. (F) Heatmap of kinase z-scores of kinases significantly enriched at at least one time point after IL-1β stimulation compared to the control samples.

Reactome pathway over-representation analysis of proteome changes clustered based on their temporal profiles revealed significant enrichment of mitochondrial respiratory electron transport (cluster 2), branched chain amino acid catabolism and fatty acid beta-oxidation (cluster 3), as well as translation of mitochondrial proteins (cluster 6) (**Figure 2B**).

Phosphoproteome profiling corroborated the global proteome findings. A total of 2,639 phosphosites were significantly regulated (adj. p-value < 0.05) 2 hours after inflammatory activation. Notably, numerous phosphorylation events characteristic for inflammatory signaling were identified already 2 hours post-IL-1β stimulation. Effects were observed on phosphosites on proteins such as IKBKE_S664^83^, MAP4K2_S328 and RELB_S37. However, the most prominent changes were observed in phosphorylation events related to cellular energy management, affecting AKT1, IRS1&2, LAMTOR1, NR4A1, RICTOR and several AMP-activated protein kinases.

The PCA indicated the greatest distance from the untreated cells 2 hours after activation, returning almost back to the baseline state 24 hours post-stimulation (**Figure 2D**). A comparison of inferred kinase activities expressed as kinase z-scores derived from a kinase-substrate enrichment analysis (KSEA), projected onto the human kinome tree is depicted in **Figure 2E** for 2h post-stimulation. KSEA results 6 and 24 hours post-stimulation are displayed in **Supplementary Figure S1** and changes in kinase z-scores throughout the total time course are summarized in **Figure 2F**. Among the most significantly early activated kinase signaling pathways were the transient activation of the MAP kinase signaling pathway, involving MAPK1, MAPK3, MAPK13 and MAPKAPK5, alongside the activation of AKT kinases (AKT1, AKT2, AKT3), inhibitor of nuclear factor kappa-B kinases (IKBKB, CHUK), and the serine/threonine kinases PRKD1 and PRKD3. Downstream targets of these pathways included ribosomal protein S6 kinases RPS6KA1 and RPS6KA3. Additionally, regulation of cell migration was indicated by STK10 and MST1R. While the global proteome levels of inflammation markers largely returned to baseline levels by 24 hours post-stimulation, kinase-substrate regulations involved in inflammatory processes remained or were later elevated (**Figure 2F, Supplementary Figure S1B**). These included the serine/threonine-protein kinases PRKG2 and PRKCB, casein kinase CSNK1D and the activin receptor ACVR2B, indicating sustained signaling and the activin receptor ACVR2B, indicating sustained signaling in specific regulatory nodes even after the primary response to inflammatory stimulation had subsided.

### Early signaling events triggered by Dexamethasone treatment show similarities to the early inflammatory signaling events in human fibroblasts

Proteome profiling identified 33 proteins that were significantly upregulated at least two-fold (log_2_FC ≥1, adj. p-value <0.05) 1 hour after treatment with Dex (**Supplementary Table S3**). Notably, almost all of these proteins were also upregulated upon inflammatory stimulation. Among the proteins induced by Dex were those typically associated with inflammation, such as NR4A1, MMP3, NFKB2 and HSD11B1 (**Figure 3A**). However, in contrast to IL-1β treatment, the inflammation markers nitric oxide synthase (NOS1), the NOS1-related extracellular matrix protein thrombospondin (THBS1), stimulator of interferon genes protein (STING1) and the inflammation modulators RIPK2 and TRAFD1 were not induced by Dex, suggesting functional specificity of these proteins. At later time points, proteins characteristic for glucocorticoid effects, such as TSC22D3^84^, FADS1 and FADS2^85^, RGCC^86^, MAOA^87^ and IL1R1^88^, exhibited significant upregulation (**Figure 3A**). Reactome pathway enrichment analysis of these proteome alterations clustered based on their temporal profiles (**Figure 3B**) showed again similarities to IL-1β treatment, particularly the induction of mitochondrial proteins (cluster 4), while also highlighting specific features of Dex treatment such as the upregulation of proteins involved in lipid metabolism (cluster 2). The PCA of the global proteome (**Figure 3C**) demonstrated transient effects along PC1, alongside more sustained changes in PC2 which persisted until 23 hours post-Dex stimulation.

**Figure 3:**
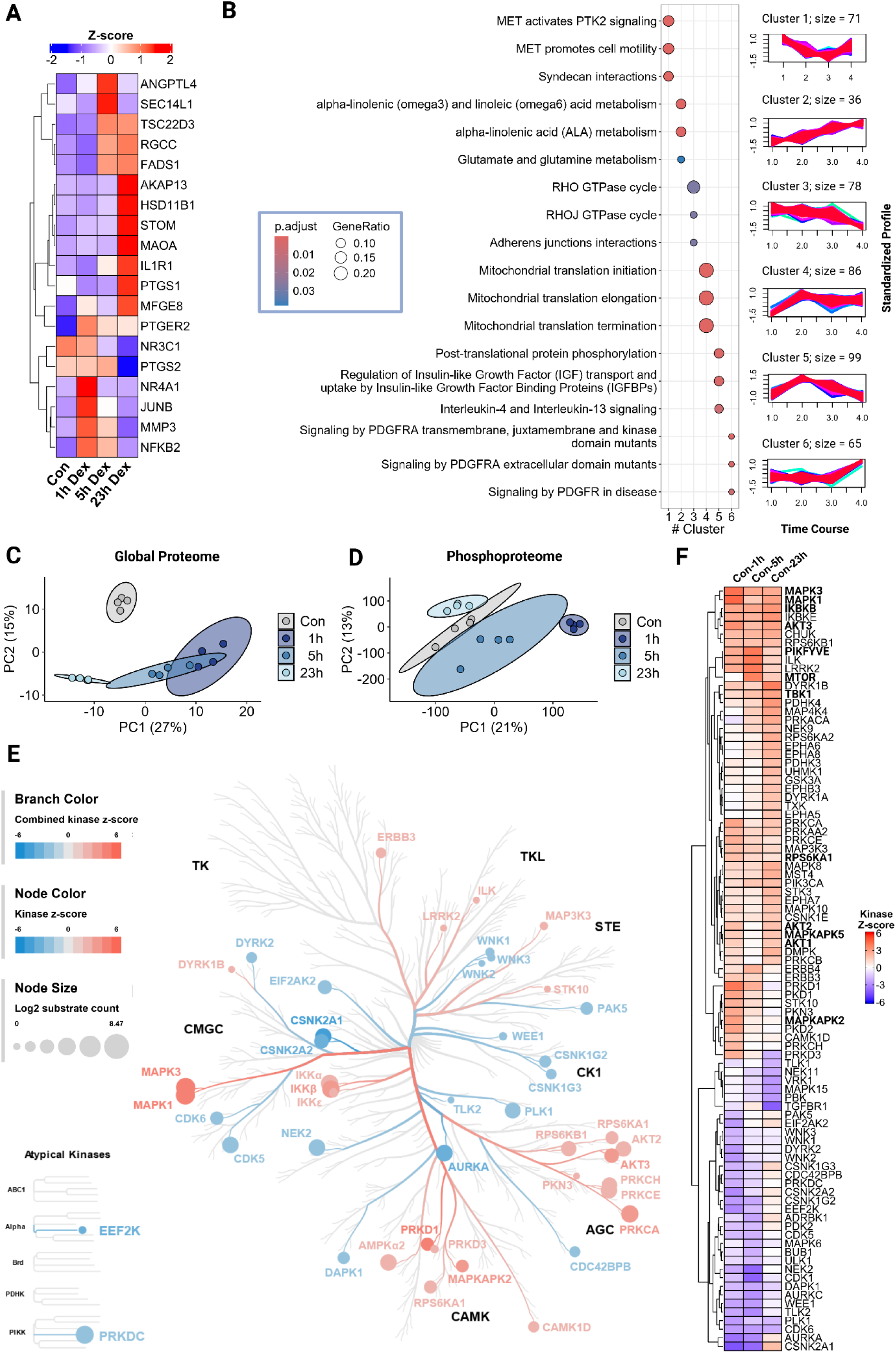
Time-resolved effects of Dexamethasone treatment on the Global Proteome and Phosphoproteome and inferred kinase activities. (A) Heatmap of z-score transformed LFQ intensities of proteins known to be altered upon Dexamethasone (Dex) treatment. Values for control samples and Dex treated samples (1, 5 and 23 h) are displayed from left to right. (B) Right panel – Clusters of proteins exhibiting similar temporal LFQ intensity profiles upon stimulation with Dex. Proteins which were significantly regulated at at least one time point compared to the control samples were subjected to the cluster analysis. Left panel – The dotplot illustrates the results of a Reactome pathway over-representation analysis performed with proteins contained in the respective clusters. The top 3 most significantly enriched pathways per cluster are shown. (C) A principal component analysis (PCA) of proteome profiles of control and Dex stimulated samples is shown. Ellipses represent 95% confidence intervals (CI). (D) A principal component analysis (PCA) of phosphoproteome profiles of control and Dex stimulated samples is shown. Ellipses represent 95% confidence intervals (CI). (E) The kinome tree depicts kinases significantly enriched for substrate phosphorylations determined by performing a kinase substrate enrichment analysis (KSEA) on class I phosphosites significantly regulated upon treatment with Dex for 1 h. Kinase z-scores indicate the statistical significance and direction of kinase activities inferred from the KSEA. (F) Heatmap of kinase z-scores of kinases significantly enriched at at least one time point after Dex stimulation compared to the control samples.

The early phosphoproteome alterations induced by Dex treatment mirrored those observed during inflammatory activation. This included phosphorylation of proteins otherwise characterized as pro-inflammatory such as IKBKE, MAP4K2 and RELB. However, Dex-specific phosphorylation events were limited, including the glucocorticoid receptor NR3C1, the phosphatidylcholine biosynthetic enzyme PCYT1A, the hormone responsive protein NDRG1 and the TGF-beta receptor signaling modulator DAB2. A comparison of kinase z-scores derived from a kinase-substrate enrichment analysis (KSEA), projected onto the human kinome tree (**Figure 3E**) revealed considerable similarities with IL-1β induced alterations (**Figure 2E**). The KSEA indicated increased activities of kinases such as MAPK1, MAPK3, AKT1, AKT2, AK3, IKBKB and MTOR, all of which are involved in cellular energy management, showing similarities to IL-1β induced effects. Interestingly, while the z-score of TBK1 decreased during inflammatory activation, it was found increased upon Dex treatment, suggesting divergent regulatory dynamics. Additionally, the dual-specificity kinase PIKFYVE, implicated in hormone-induced signaling, was specifically activated in response to Dex.

To comprehensively assess the proportion of shared and specific effects of IL-1β and Dex, we performed a systematic comparison of the proteome and phosphoproteome alterations (**Figure 4)**. At the first timepoint of the temporal profiles (2 hours for IL-1β, 1 hour for Dex), 67% and 72% of significantly upregulated proteins (adj. P-value ≤0.05, fold change ≥1) upon IL-1β and Dex treatment, respectively, were also significantly upregulated in the other condition (**Figure 4A)**. A similar trend was observed for significantly downregulated proteins (adj. P-value ≤0.05, fold change ≥-1), with 52% and 70% shared regulations, respectively. With increasing post-treatment time, the proportion of shared regulated proteins decreased to 50% and 61% for upregulated proteins and to 42% and 67% for downregulated proteins, for IL-1β and Dex treatment, respectively. Since the number of significantly regulated proteins drastically decreased between 6 and 24 hours post IL-1β treatment, the majority of them was shared with altered proteins 23 hours post-Dex treatment, while nearly all of the alterations at this late time point were Dex-exclusive. Interestingly, while the number of significantly altered proteins upon IL-1β stimulation continuously decreased over the time, Dex-induced regulations decreased from 1 hour to 5 hours, before increasing again between 5 hours and 23 hours. Similar trends were observed on the phosphoproteome level represented by significantly enriched kinases (P-value ≤0.05) derived from the KSEA based on significantly regulated phosphosites in each condition (**Figure 4B**). The number of kinases was similar at the early time points and also decreased to a similar extent until the middle-term time points, for both treatments, indicated by the right-hand barplot in **Figure 4B**. However, just like observed on proteome level, while the proportion of IL-1β altered kinases further decreased until 24 hours post-treatment, the number of Dex-perturbed kinases was enhanced again at 23 hours. Remarkably, more than half of the sig. regulated kinases 23 hours post-Dex treatment, 25 out of 47, were exclusively dysregulated upon Dex treatment at this specific time point, indicating a comparably late onset of a broad panel of Dex-exclusive effects.

**Figure 4:**
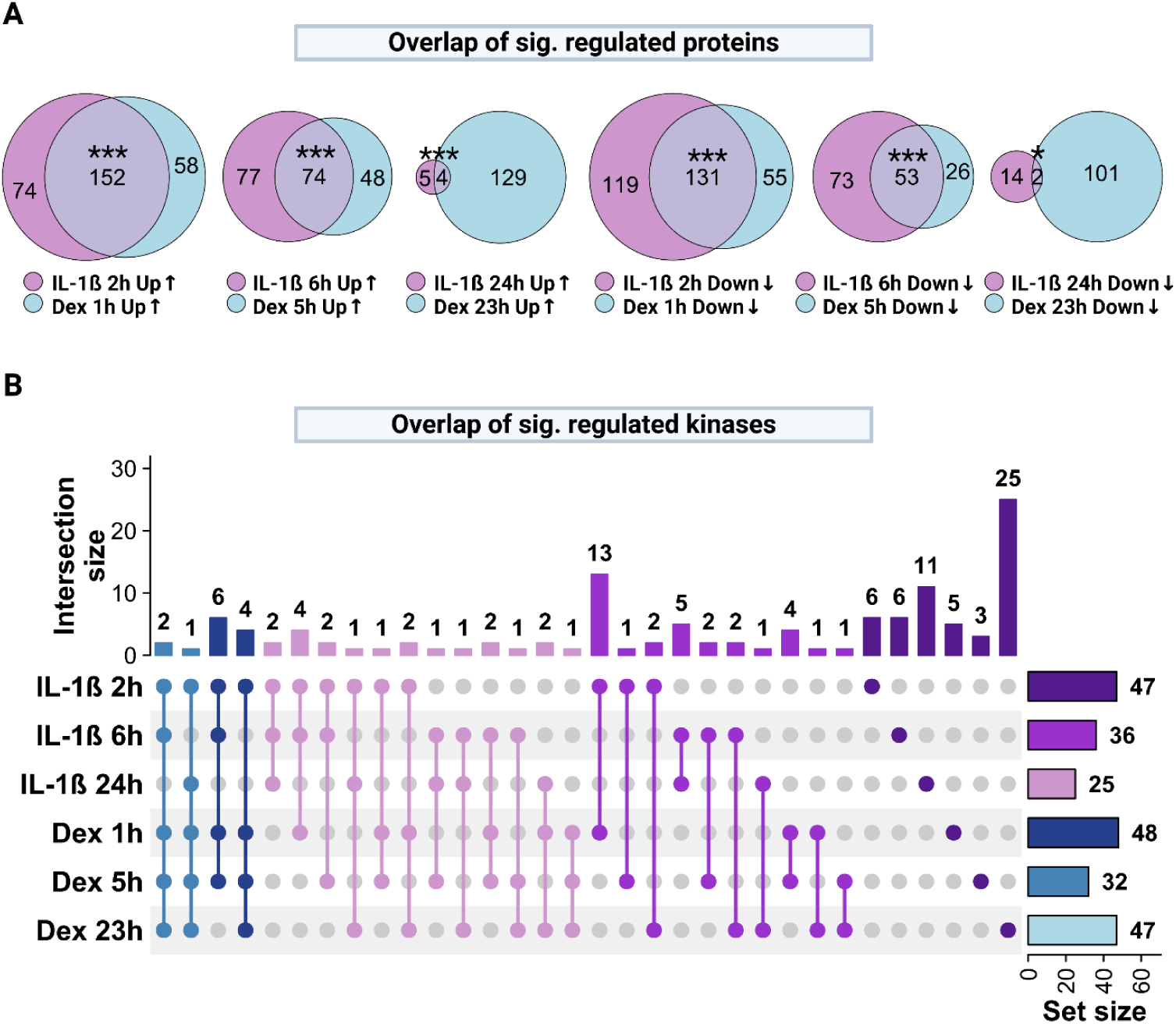
Shared and specific proteomic and phosphoproteomic effects of IL-1β and Dexamethasone treatment. (A) The Venn diagrams illustrate shared and unique significantly up-(top panel) or down-regulated (bottom panel) proteins (adj. *P*-value ≤0.05, fold change ≥1 or ≤-1) upon IL-1β (purple) and Dexamethasone treatment (blue) in comparison to control samples. Diagrams for the matched short-, middle- and long-term treated samples are shown from left to right. Statistical significance was calculated using a hypergeometric test (***P-value <0.001, *P-value <0.05,). (B) The UpSet plot illustrates shared and unique significantly regulated kinases derived from the kinase substrate enrichment analysis in comparison to control samples. The top bar plot demonstrates the size of the intersection between the respective sets illustrated via the lines and dots in the rows below. The bar plot on the right side shows the total set size of the sig. regulated kinases per treatment and time point.

To obtain a more detailed view on the molecular events induced by IL-1β and Dex treatment and their underlying regulatory mechanisms, we integrated the proteomic and phosphoproteomic datasets into bilevel causal perturbation networks using CausalPath^53^ (**Figure 5, Supplementary Figure S4 A and B**). The analysis revealed central molecular hubs in the early response network 2 hours post-IL-1β stimulation (**Figure 5A**), which were indicated by a large number of causal edges connecting the respective nodes (proteins). These hubs included AKT1, CREBBP, STAT1, STAT3, MAPK1, NFKB1, CDKN1A and PRKCA. Similarly, AKT1, NFKB1, PRKCA and CREBBP were also identified as central protein hubs in the network obtained 1 hour post-Dex treatment (**Figure 5B**). Upon this short-term Dex treatment, more connections to the stress-activated serine/threonine-protein kinase MAPKAPK2, and less connections to STAT1, STAT3 and MAPK1 were obtained (**Figure 5B**). Despite similarities, the causal network established upon 2 hours post-IL-1β treatment in general yielded more causal conjectures than the network based on the perturbations observed 1 hour post-Dex treatment. Per definition, a causal conjecture is defined as the pairing of a causal knowledge with supporting experimental data, which together declare that “one molecular change is the cause of another molecular change”^53^.

**Figure 5:**
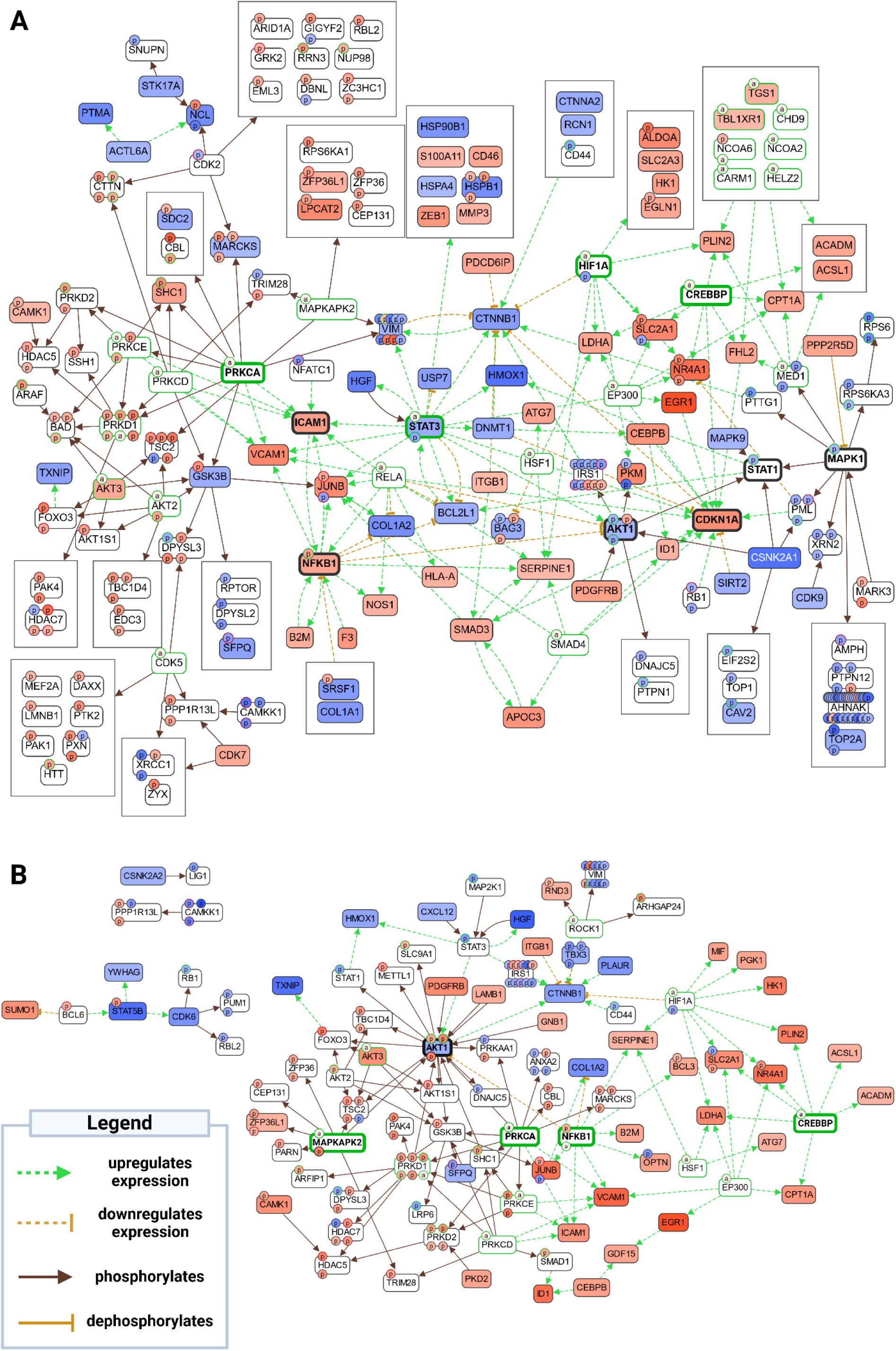
Bilevel CausalPath analysis using global proteomic and phosphoproteomic data. (A) IL-1β 2 h vs. Con (B) Dex 1h vs. Con – Red indicates up-regulated total protein quantity (rectangular boxes) or protein feature quantity (Circles). Blue indicates total protein quantity (rectangular boxes) or protein feature quantity (Circles). White colored proteins with colored protein features indicate either no change in quantity of the unphosphorylated protein or missing quantitative data at unphosphorylated protein level. Continuous-line brown arrows indicate phosphorylation events. Dashed green lines represent a positive correlation of gene expression changes, whereas dashed orange lines indicate a negative correlation of gene expression changes. Phosphosites with a green frame are known to be activating, phosphosites with a red frame are known to be deactivating. Gene expression products marked with “a” are annotated by the CausalPath algorithm due to the concerted upregulation of associated gene expression products.

### Dexamethasone inhibits pro-inflammatory signaling but elicits distinct signaling events

In order to investigate the molecular consequences of Dex treatment affecting already inflammatory activated cells, the similar time course analysis was applied to cells first stimulated with IL-1β followed by Dex treatment one hour thereafter. Consequently, the molecular profiles of the IL-1β treated cells were used as reference here, resulting in matched post-treatment time points for all comparisons. A PCA of the proteomes obtained upon this combined treatment and upon IL-1β treatment only (**Figure 6A, left panel)** demonstrated that the difference was smallest at 2 hours total incubation time per each condition and greatest at 24 hours total incubation time per each condition. In contrast, in the PCA of the corresponding phosphoproteomes, the small, but notable group difference observed after 2 hours completely disappeared after 24 hours (**Figure 6A, right panel**). Compared to sole IL-1β stimulation, the additional subsequent Dex treatment induced the expression of anti-inflammatory proteins including MFGE8, HSD11B1 and TSC22D3 (**Figure 6B**). In a similar notion, the expression of pro-inflammatory proteins, such as PTGS2, SASH1 and IL33 was decreased (**Figure 6B**). Notably, as also reflected by the PCA of the global proteomes, the majority of the proteins significantly regulated upon additional Dex treatment exhibited the greatest contrast at 24 hours post-IL-1β stimulation. As observed in the temporal proteome profile upon sole Dex treatment, this again points towards specific Dex-effects mainly taking place only after long term (23 or 24 hours) treatment.

**Figure 6:**
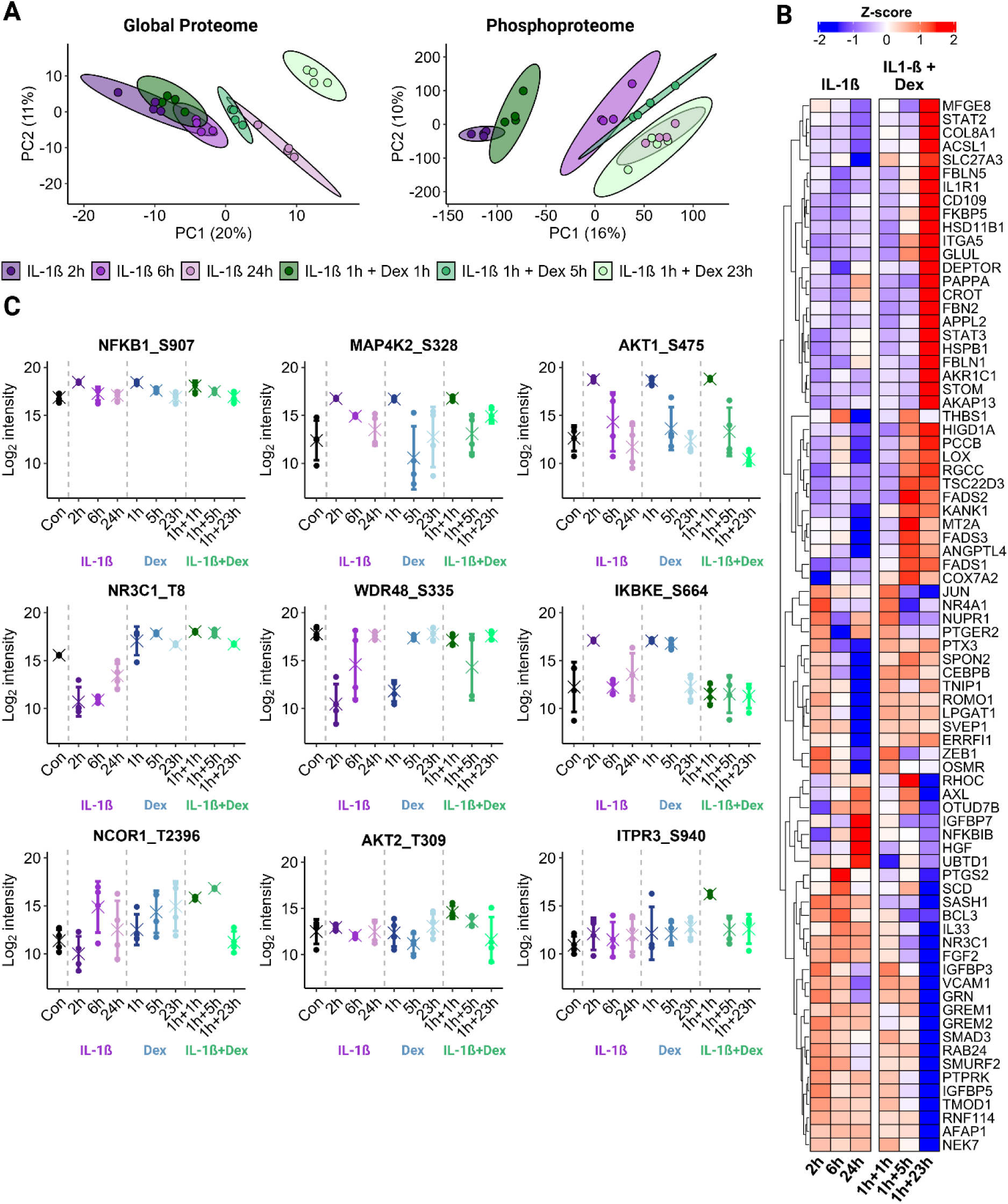
Time-resolved effects of a combined IL-1β and Dexamethasone treatment on the Global Proteome and Phosphoproteome and inferred kinase activities. (A) *Left Panel.* Global Proteome PCA *Right Panel.* Phosphoproteome PCA (C) Heatmap of LFQ intensities of selected proteins sig. regulated at at least one time point in the comparison IL-1β + Dex vs. IL-1β

The phosphorylation events observed upon combined treatment showed similarities to the single treatments, but also displayed specific properties. Representatively, phosphorylation of NFKB1 at serine-907, MAP4K2 at serine-328 and AKT1 at serine-475 showed similar temporal profiles in all three conditions (**Figure 6C**, top row). Conversely, phosphorylation of the glucocorticoid receptor NR3C1 at threonine-8 was decreased upon inflammatory stimulation, but increased by Dex and the combined treatment, confirming effective Dex treatment in both cases. Interestingly, phosphorylation of WD repeat-containing protein (WDR48), a regulatory protein playing a crucial role in several cellular processes by modulating protein ubiquitination, at serine-335, was decreased upon both individual short-term treatments, but not upon combined treatment (**Figure 6C**, middle row - middle panel). In similar notion, IKBKE phosphorylation at serine-664 was induced upon both individual short-term stimulations, but not upon combined treatment (**Figure 6C**, middle row - right panel). While such observations may account for IL-1β- or Dex-specific signaling, some events specific for the combined treatment were observed as well. This applied e.g. to enhanced phosphorylation of the nuclear receptor corepressor 1 (NCOR) at threonine-2396. Enhanced phosphorylation upon the combined treatment, but not of the individual treatments, was also observed on serine-940 at ITPR3, a calcium channel protein located on the endoplasmic reticulum.

### Lysolipid and oxylipin analysis indicate involvement of lipid metabolism for energy management

While whole cell lysates were used for proteomics and phosphoproteomics analyses, cell supernatants were analyzed with regard to lysolipids and oxylipins. Compared to the control condition, IL-1β induced the formation of numerous lysolipids, mainly lysophosphatidylcholines (LPCs) and lysophosphatidylethanolamines (LPEs), 2 hours and 6 hours post-treatment, demonstrating the activity of phospholipases upon inflammatory stimulation (**Figure 7A**). The oxylipins PGA3, 15deoxy-PGJ2, LTB4, 5-HEPE and 5-HETE, amongst several other, were also found induced in a similar temporal behavior (**Supplementary Table S4**). Several unsaturated fatty acids such as AA, DGLA and DPA were found increased, whereas EPA, linolenic acid and stearidonic acid were found decreased. Oxylipins with pro-resolving properties such as 14(15)-EET were found decreased, others such as Resolvin E2 were found increased. In general, the overall effects of IL-1β on the lipid composition of the cell supernatant attenuated 6 hours post-treatment. Upon 24 hours post-IL-1β stimulation, the effects had completely resolved.

**Figure 7:**
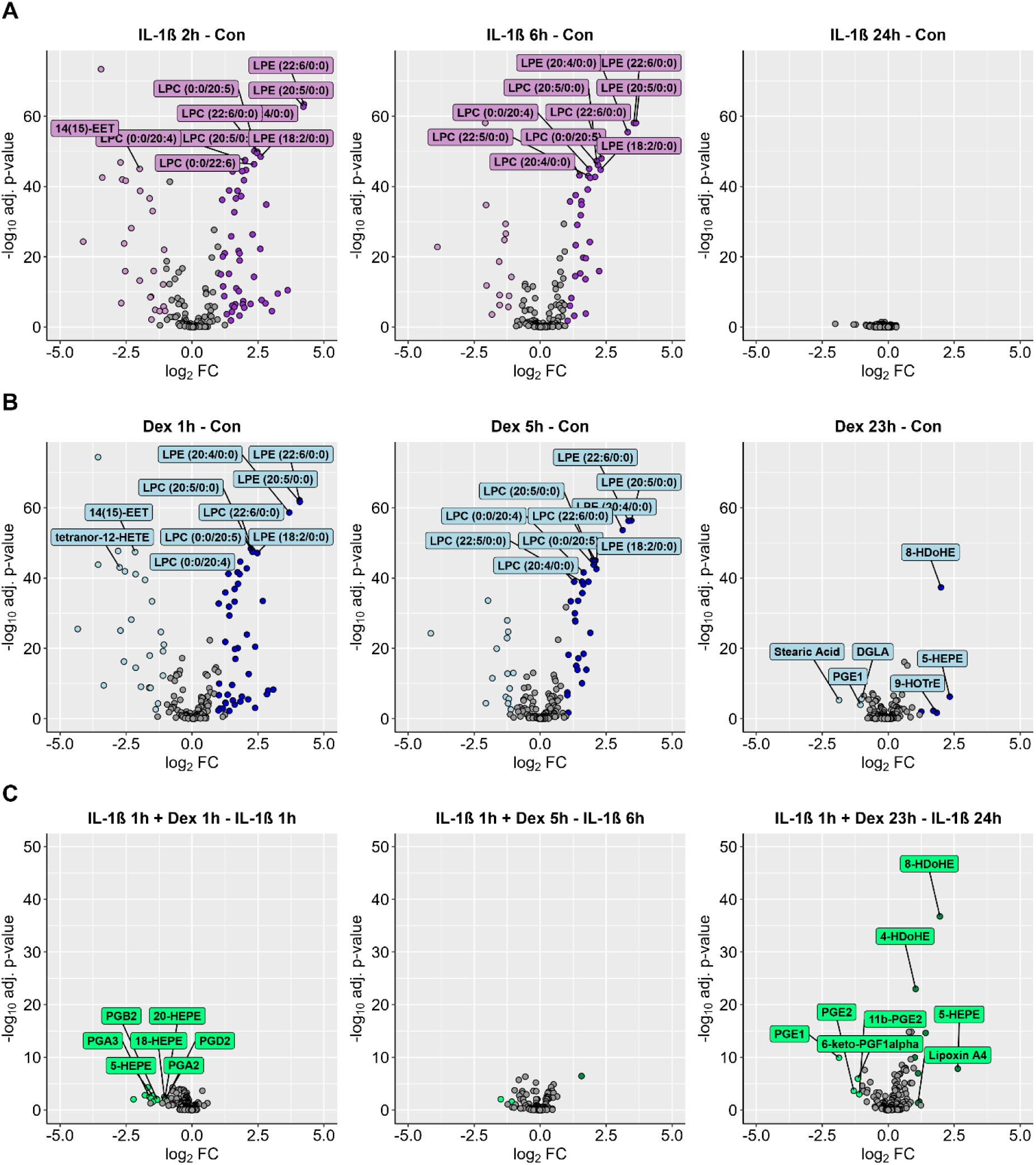
Oxylipin and fatty acid data. Volcano plots showing the (A) IL-1β-(B) Dex- and (C) IL-1β + Dex-induced oxylipin and fatty acid perturbation profiles of Detroit-551 cell supernatants. Significant regulations after multiple testing correction are colored (adj. *P*-value ≤0.05, fold change ≥1 or ≤-1). The up to top 10 significant regulations with the lowest adj. p-values are labelled, excluding isoforms and non-explicitly defined molecules.

The effects of Dex treatment on the oxylipins and fatty acids were again highly similar to those of IL-1β treatment, including regulation of numerous LPCs and LPEs upon short- and middle-term treatment (**Figure 7B**). However, in contrast to IL-1β treatment, 5-HEPE, 8-HDoHE, 9-HOTrE and were found still upregulated as well as DGLA, Stearic acid and PGE1 downregulated after 23 hours. As observed in the proteome and phosphoproteome upon Dex-treatment, this indicates Dex-specific events upon long-term incubation.

An effect of Dex on the oxylipin and fatty acid composition of the supernatants of IL-1β pre-treated cells was already detectable after 1 hour (**Figure 7C**) and was most pronounced after 24 hours. While the prostaglandin PGE2 was found downregulated at the latest time point, the pro-resolving oxylipins 8-HDoHE and 5-HEPE were found up-regulated.

### A global view on signaling events associated with an inflammatory response demonstrates the complexity of cellular resource governance

The presented study provides insights into the proteomic alterations, the dynamic signaling and the metabolic responses of human embryonic fibroblasts to inflammatory stimulation (IL-1β) and anti-inflammatory treatment (Dexamethasone), utilizing an integrated proteomics, phosphoproteomics and oxylipin/fatty acid analysis approach.

Our results revealed that both IL-1β and Dex trigger strikingly similar early proteome, phosphoproteome and oxylipin/fatty acid alterations, while Dex-specific effects are mainly observed after prolonged treatment. Thus, the data suggests that similar signaling events are triggered by two different kinds of treatment. These early signaling events, particularly those related to cell growth, adhesion, migration, and metabolism, suggest a greater emphasis on cellular resource governance rather than a purely inflammatory or anti-inflammatory response.

### Cellular resource governance and stress management in inflammation

A key observation in this study was the prominent early involvement of metabolic pathways, including mitochondrial function, β-oxidation, and glycolysis, in response to both IL-1β and Dex treatments. These processes likely serve to meet the increased energy demands necessary for transcriptional and translational activation during the initial stress response.

The data also suggested that increased mitochondrial electron transport chain activity and ATP production might contribute to higher reactive oxygen species (ROS) generation^89^ causing cell stress, known features of chronic inflammation. It can thus be hypothesized that the role of FOXO3 mediating stress adaptation and extending life span^90^ may be linked to the cellular requirement to get rid of dysfunctional proteins and organelles *via* autophagy after such a challenge. Delayed disposal may cause secondary inflammatory responses, consequently contributing to chronic inflammation, “inflamm-aging” and associated diseases affecting the life expectancy.

Only after successful management of these challenges, the specific consequences of the different kinds of activation seem to become prevalent. This interpretation may suggest some important consequences. The difference between “mediator” and “indicator” of processes needs to become more precisive. To give an example, acute-phase proteins known to be induced upon inflammation represent indicators of inflammation rather than mediators. Indeed, increased levels of acute-phase proteins such as c-reactive protein indicate the presence of inflammation, while promoting the resolution of inflammation^91^.

### Reassessing kinase activity in inflammation

The involvement of kinase activation, particularly NF-kappa B, STAT, and MAPK pathways, is a central theme upon both, the IL-1β and Dex treatments. Our findings indicate that the involvement of kinase activation in course of inflammatory or anti-inflammatory stimulation seems to be rather complex. These findings evidence that kinase activation observed upon inflammatory stimulation does not necessarily represent a pro-inflammatory event. Instead, these kinase activations may play logistical roles that are necessary to establish the inflammatory environment, such as energy regulation, without necessarily promoting the inflammation itself. This is evident from the Dex-treated fibroblasts, where kinase activation was observed in pathways often associated with pro-inflammatory processes, despite the overall anti-inflammatory outcome of the treatment. In support of this interpretation, NF-kappa B signaling independent of any inflammation was identified as a critical functional factor for the mammalian nervous system.^92^ This raises important questions regarding the classification of kinases as “pro-inflammatory” or “anti-inflammatory”. Instead, our data suggest a need for a more nuanced understanding where kinases may act as facilitators of both inflammation and resolution, depending on the cellular context and the balance of metabolic and signaling demands.

The analysis of the effects of dexamethasone on the background of pre-triggered inflammatory response verified the known inhibitory functions of Dex and resulted in the expression of anti-inflammatory proteins and oxylipins, but hardly affected the signaling events involving MAP and AKT kinases.

### Mitochondrial fitness as potential contributor to the temporal effectiveness of resolution of inflammation

An additional finding of this study is the difference in the temporal dynamics of the inflammatory response between embryonic and adult fibroblasts. In contrast to our previous work^16^ using adult fibroblasts, the embryonic cells employed in this study displayed much faster resolution of the inflammation, with protein markers such as COX2 (PTGES2), NF-kappa-B p100, and JUN being rapidly induced within 2 hours, but subsequently returning to baseline levels at 24 hours. This accelerated response mirrors findings in *in vivo* models of tissue injury, where fetal organisms resolve inflammation more quickly than their adult counterparts.^93^

The underlying cause of this enhanced inflammatory resolution could lie in a more robust mitochondrial function and more efficient regeneration of damaged mitochondria in embryonic fibroblasts^91^, as mitochondrial dysfunction has been implicated in the chronicity of inflammation in aging tissues^94^ and has been described to be associated with long-term cell culture^95^. These findings underscore the importance of mitochondrial health and regenerative capacity in controlling the duration and severity of inflammation and raise the possibility that targeting mitochondrial function could be a viable strategy for treating chronic inflammatory conditions in aging or metabolically compromised tissues.

## CONCLUSION

In summary, this study offers a temporally resolved, multi-omics-driven analysis of human embryonic fibroblasts’ response to inflammatory and anti-inflammatory stimuli. The early signaling events triggered by IL-1β and Dexamethasone converge on metabolic pathways essential for resource governance, suggesting that cellular energy management is an integral part of the inflammatory response. These findings challenge the traditional view of kinase activity in inflammation, highlighting the need for a more refined classification of kinase functions. The comparison of embryonic and adult fibroblasts reveals insights into the eventual role of mitochondrial function in resolving inflammation, potentially relevant for better understanding processes underlying chronic inflammation.

Our findings highlight the complexity of fibroblast-mediated inflammation and underscore the importance of metabolic regulation in both promoting and resolving inflammatory responses. While the methodology employed here provides a comprehensive view of the early perturbations, future studies could benefit from expanding these time-course analyses to further elucidate the molecular underpinnings of inflammation and its resolution.

These insights may contribute to the knowledge base relevant for therapeutic strategies aimed at improving mitochondrial function and selectively targeting signaling pathways involved in chronic inflammatory diseases.

## Acknowledgements

Figures were created with in BioRender.com.

## Funding and additional information

Work at The Novo Nordisk Foundation Center for Protein Research (CPR) is funded in part by a donation from the Novo Nordisk Foundation (NNF14CC0001). This work has also been funded as part of EPIC-XS project under the grant agreement no. 823839 funded by the Horizon 2020 programme of the European Union. P.B. was supported by the “International Exchange Program” (Vienna Doctoral School in Chemistry, DoSChem), the funding program “Internationale Kommunikation” (Österreichische Forschungsgemeinschaft, ÖFG) and the “Erasmus+ Traineeship Mobility” program.

## Autor contributions

C.G. and P.J.-B. conceptualization; P.J.-B., A.M.V., G.H. and L.S. methodology; P.J.-B. and G.H. investigation; P.J.-B. formal analysis; C.G. and P.B. writing – original draft; C.G, P.B, A.M.-V., G.H., L.S., J.V.O. writing – review and editing.

