## Supplementary Figure for "Phosphoproteomics highlights complex resource management upon inflammatory stimulation of fibroblasts"

### SUPPLEMENTARY FIGURES

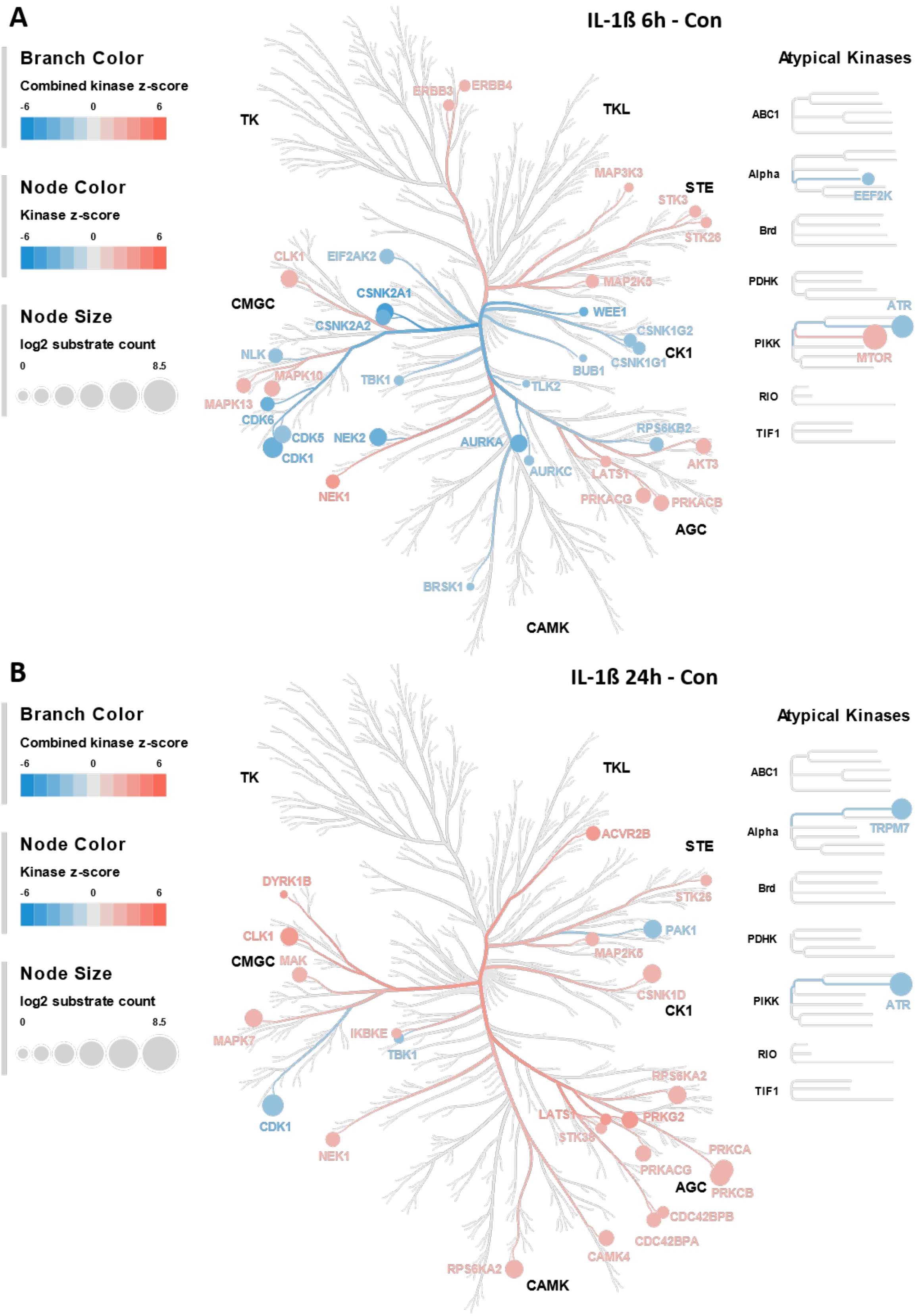


**Supplementary Figure S1: Time-resolved effects of IL-1ß on the Global Proteome and Phosphoproteome and inferred kinase activities.** The kinome trees depict kinases significantly enriched for substrate phosphorylations determined by performing a kinase substrate enrichment analysis (KSEA) on class I phosphosites significantly regulated upon treatment with (A) IL-1ß for 6 h and (B) IL-1ß for 24 h, each compared against control samples.

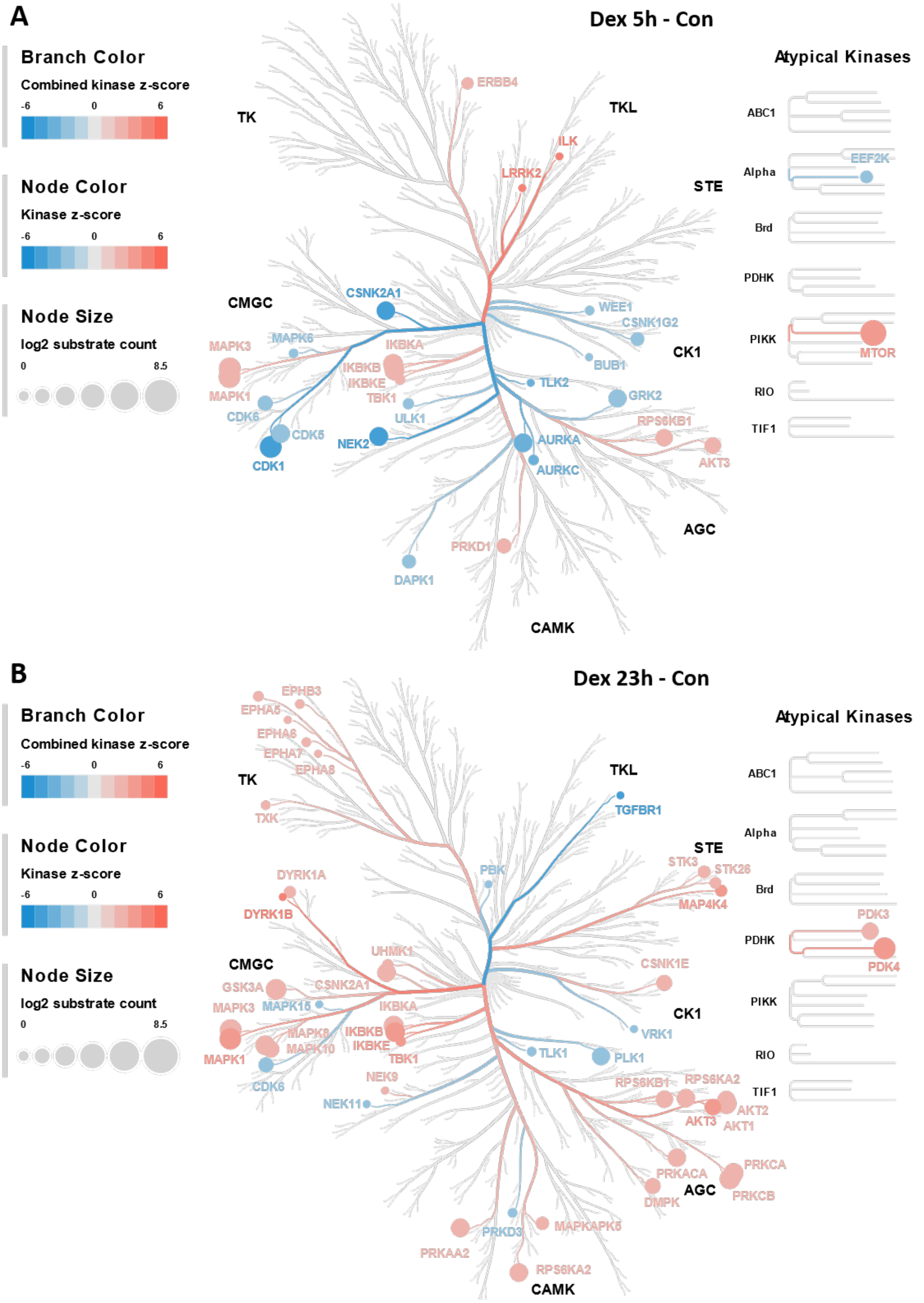


**Supplementary Figure S2: Time-resolved effects of Dexamethasone on the Global Proteome and Phosphoproteome and inferred kinase activities.** The kinome trees depict kinases significantly enriched for substrate phosphorylations determined by performing a kinase substrate enrichment analysis (KSEA) on class I phosphosites significantly regulated upon treatment with (A) Dexamethasone for 5 h and (B) Dexamethasone for 23 h, each compared against control samples.

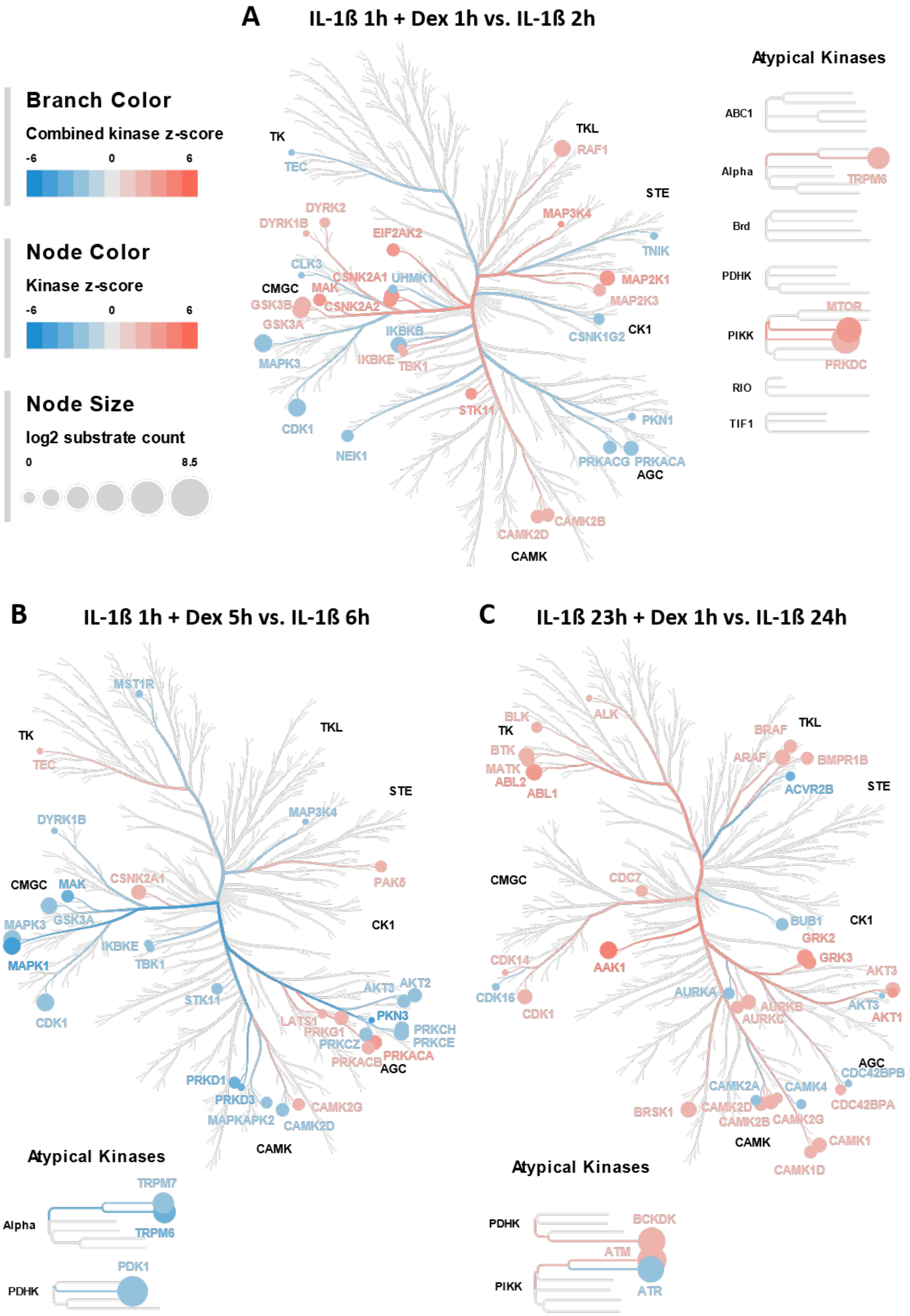


**Supplementary Figure S3: Time-resolved effects of the combined IL-1ß + Dexamethasone (Dex) treatment on the Global Proteome and Phosphoproteome and inferred kinase activities in comparison to IL-1ß treatment.** The kinome trees depict kinases significantly enriched for substrate phosphorylations determined by performing a kinase substrate enrichment analysis (KSEA) on class I phosphosites significantly regulated upon treatment with (A) IL-1ß for 1 h + Dex for 1 h compared against treatment with IL-1ß for 2 h (B) IL-1ß for 1 h + Dex for 5 h compared against treatment with IL-1ß for 6 h (C) IL-1ß for 1 h + Dex for 23 h compared against treatment with IL-1ß for 24 h.

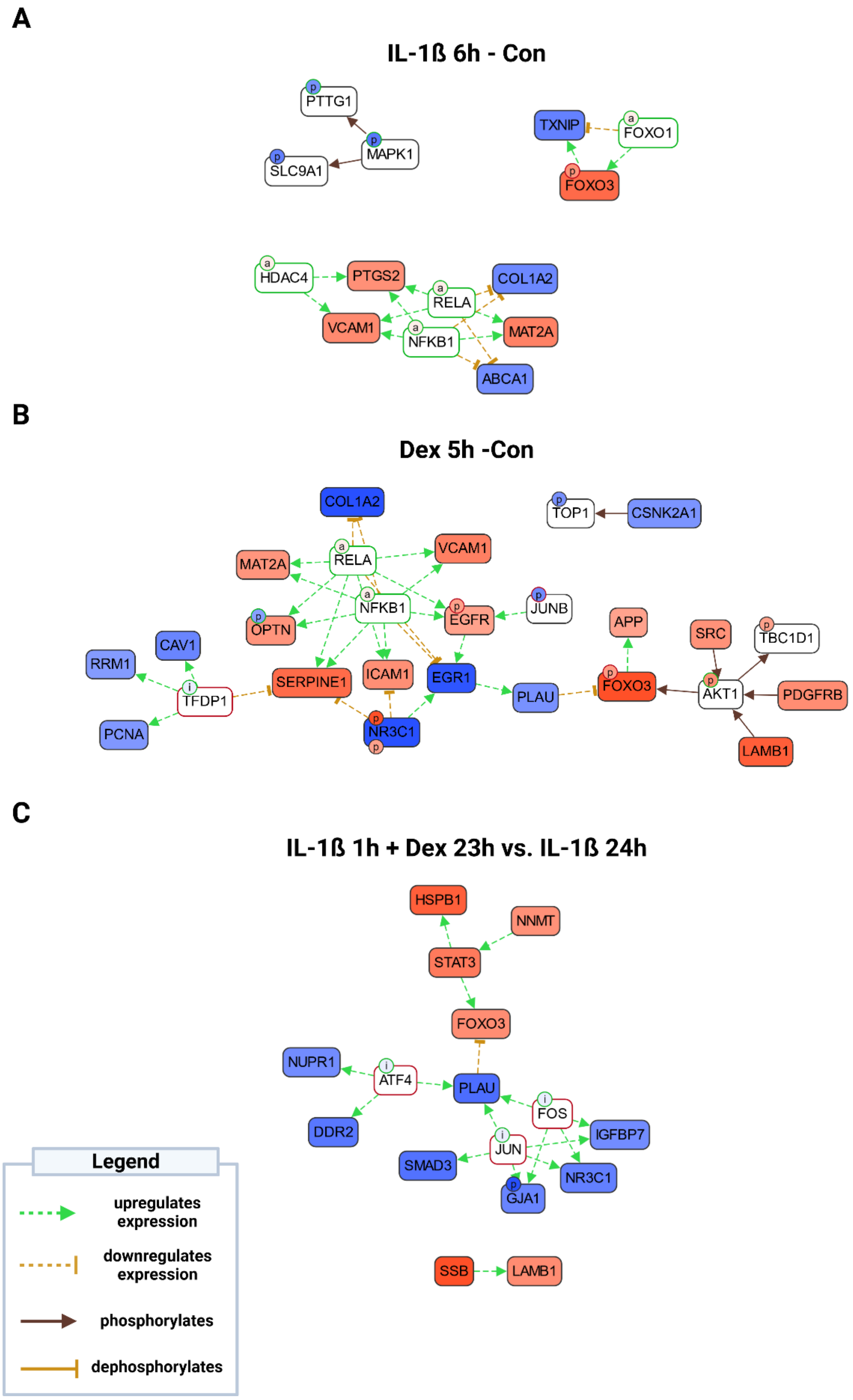


**Supplementary Figure S4: Bilevel CausalPath analysis using global proteomic and phosphoproteomic data.** (A) IL-1ß 6 h vs. Con (B) Dex 5h vs. Con (C) IL-1ß 1 h + Dex 23 h vs. IL-1ß

24 h – Red indicates up-regulated total protein quantity (rectangular boxes) or protein feature quantity (Circles). Blue indicates total protein quantity (rectangular boxes) or protein feature quantity (Circles). White colored proteins with colored protein features indicate either no change in quantity of the unphosphorylated protein or missing quantitative data at unphosphorylated protein level. Continuous-line brown arrows indicate phosphorylation events. Dashed green lines represent a positive correlation of gene expression changes, whereas dashed orange lines indicate a negative correlation of gene expression changes. Phosphosites with a green frame are known to be activating, phosphosites with a red frame are known to be deactivating. Gene expression products marked with “a” are annotated by the CausalPath algorithm due to the concerted upregulation of associated gene expression products.

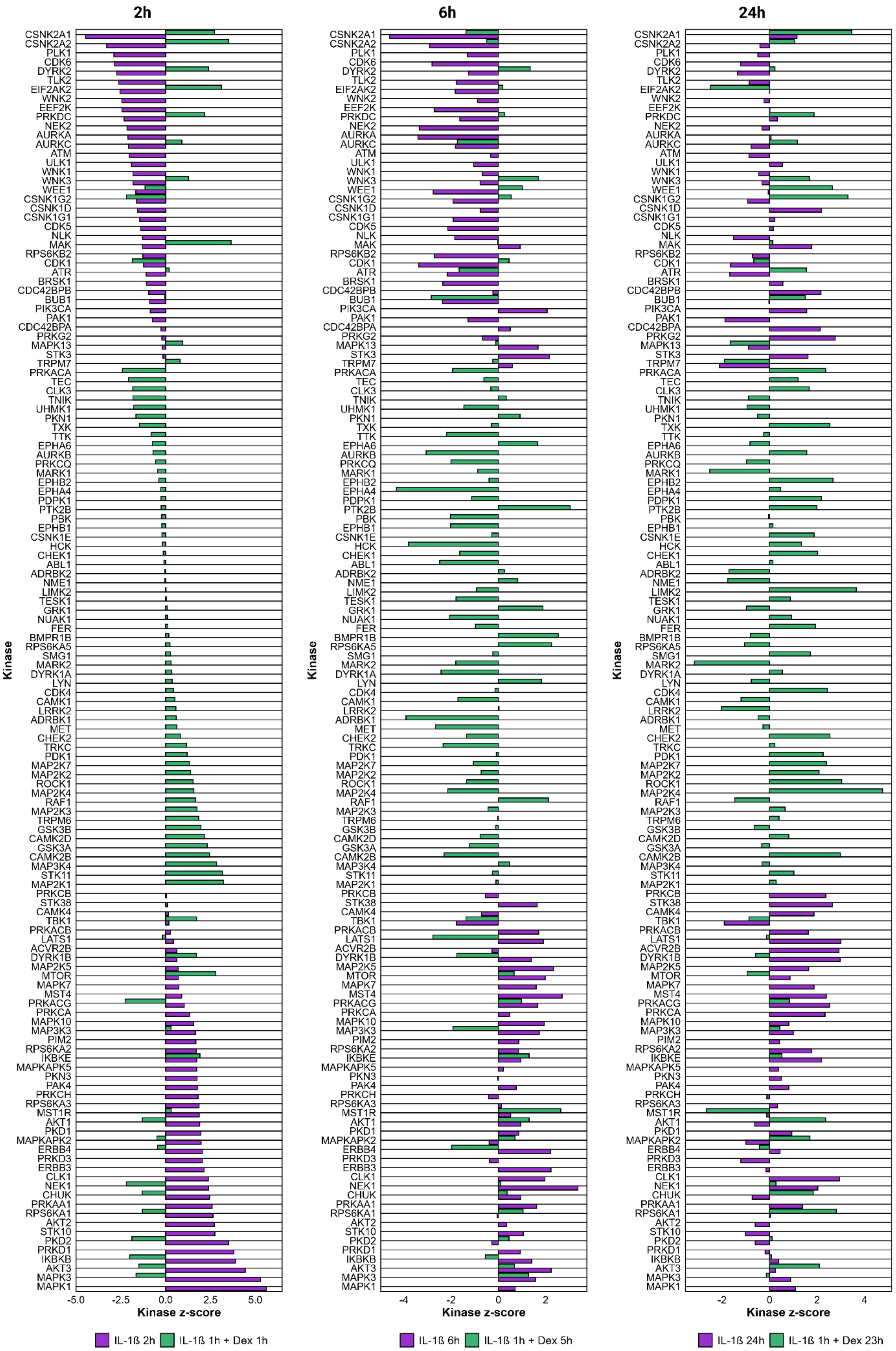
2h
6h
24h
Kinase
CSNK2A1
CSNK2A2
PLK1
CDK6
DYRK2
TLK2
EIF2AK2
WNK2
EEF2K
PRKDC
NEK2
AURKA
AURKC
ATM
ULK1
WNK1
WNK3
WEE1
CSNK1G2
CSNK1D
CSNK1G1
CDK5
NLK
MAK
RPS6KB2
CDK1
ATR
BRSK1
CDC42BPB
BUB1
PIK3CA
PAK1
CDC42BPA
PRKG2
MAPK13
STK3
TRPM7
PRKACA
TEC
CLK3
TNIK
UHMK1
PKN1
TXK
TTK
EPHA6
AURKB
PRKCQ
MARK1
EPHB2
EPHA4
PDPK1
PTK2B
PBK
EPHB1
CSNK1E
HCK
CHEK1
ABL1
ADRBK2
NME1
LIMK2
TESK1
GRK1
NUAK1
FER
BMPR1B
RPS6KA5
SMG1
MARK2
DYRK1A
LYN
CDK4
CAMK1
LRRK2
ADRBK1
MET
CHEK2
TRKC
PDK1
MAP2K7
MAP2K2
ROCK1
MAP2K4
RAF1
MAP2K3
TRPM6
GSK3B
CAMK2D
GSK3A
CAMK2B
MAP3K4
STK11
MAP2K1
PRKCB
STK38
CAMK4
TBK1
PRKACB
LATS1
ACVR2B
DYRK1B
MAP2K5
MTOR
MAPK7
MST4
PRKACG
PRKCA
MAPK10
MAP3K3
PIM2
RPS6KA2
IKBKE
MAPKAPK5
PKN3
PAK4
PRKCH
RPS6KA3
MST1R
AKT1
PKD1
MAPKAPK2
ERBB4
PRKD3
ERBB3
CLK1
NEK1
CHUK
PRKAA1
RPS6KA1
AKT2
STK10
PKD2
PRKD1
IKBKB
AKT3
MAPK3
MAPK1
-5.0 -2.5 0.0 2.5 5.0
-4 -2 0 2
-2 0 2 4
Kinase z-score
IL-1ß 2h
IL-1ß 1h + Dex 1h
IL-1ß 6h
IL-1ß 1h + Dex 5h
IL-1ß 24h
IL-1ß 1h + Dex 23h

**Supplementary Figure S5:** Bar plots of kinase z-scores from kinase substrate enrichment analysis, grouped per total treatment duration for IL-1ß (purple) and IL-1ß + Dex (green) treated samples.

### SUPPLEMENTARY TABLES

**Table S1:** List of spiked internal eicosanoid standards and their respective final concentration in each sample.

| Internal standard | Abbreviation | c [pg/µL] |
|---|---|---|
| 12S-hydroxyeicosatetraenoic acid-d8 | 12S-HETE-d8 | 6.67 |
| 15S-hydroxyeicosatetraenoic acid-d8 | 15S-HETE-d8 | 6.67 |
| 5-oxo-eicosatetraenoic acid-d7 | 5-OxoETE-d7 | 20.00 |
| 11,12-dihydroxy-5Z,8Z,14Z-eicosatrienoic acid-d11 | 11,12-DiHETrE-d11 | 6.67 |
| prostaglandin E2-d4 | PGE2-d4 | 13.33 |
| 20-hydroxyeicosatetraenoic acid-d6 | 20-HETE-d6 | 6.67 |

**Table S2:** Inclusion list of 33 precursor masses of eicosanoids and fatty acids utilized for mass spectrometric analysis.

| [m/z] | | |
|---|---|---|
| 254.2245 | 313.2384 | 337.2384 |
| 275.2011 | 315.1966 | 343.2279 |
| 277.2167 | 317.2122 | 348.3069 |
| 279.2324 | 319.2279 | 349.2020 |
| 281.2480 | 321.2435 | 351.2177 |
| 283.2637 | 325.2382 | 353.2328 |
| 293.2122 | 327.2324 | 355.2428 |
| 295.2279 | 327.2781 | 357.2585 |
| 301.2168 | 329.2480 | 359.2222 |
| 303.2324 | 333.2071 | 367.3576 |
| 311.2228 | 335.2222 | 375.2171 |

**Table S3:** Proteins significantly upregulated more than two-fold upon 1h Dexamethasone treatment, compared against control samples.

| Gene | UniProt ID | Protein Description | $log_2FC$ | adj. p-value |
|---|---|---|---|---|
| EGR1 | P18146 | Early growth response protein 1 | 3.40 | 0.00 |
| PTGER2 | P43116 | Prostaglandin E2 receptor EP2 subtype | 2.89 | 0.00 |
| NR4A1 | P22736 | Nuclear receptor subfamily 4 group A member 1 | 2.78 | 0.00 |
| CHKB | Q9Y259 | Choline/ethanolamine kinase | 2.77 | 0.01 |
| VCAM1 | P19320 | Vascular cell adhesion protein 1 | 2.71 | 0.00 |
| RAB24 | Q969Q5 | Ras-related protein Rab-24 | 2.49 | 0.03 |
| MMP3 | P08254 | Stromelysin-1 | 2.35 | 0.00 |
| SLC39A8 | Q9C0K1 | Metal cation symporter ZIP8 | 1.93 | 0.00 |
| JUNB | P17275 | Transcription factor JunB | 1.84 | 0.00 |
| FOXO3 | O43524 | Forkhead box protein O3 | 1.83 | 0.00 |
| CEMIP | Q8WUJ3 | Cell migration-inducing and hyaluronan-binding protein | 1.79 | 0.00 |
| SRD5A3 | Q9H8P0 | Polyprenol reductase | 1.77 | 0.03 |
| RB1CC1 | Q8TDY2 | RB1-inducible coiled-coil protein 1 | 1.66 | 0.00 |
| POTEKP | Q9BYX7 | Putative beta-actin-like protein 3 | 1.61 | 0.01 |
| CA12 | O43570 | Carbonic anhydrase 12 | 1.58 | 0.00 |
| CCN2 | P29279 | CCN family member 2 | 1.49 | 0.00 |
| UIMC1 | Q96RL1 | BRCA1-A complex subunit RAP80 | 1.41 | 0.03 |
| ERRFI1 | Q9UJM3 | ERBB receptor feedback inhibitor 1 | 1.41 | 0.00 |
| ZNF703 | Q9H7S9 | Zinc finger protein 703 | 1.38 | 0.00 |
| GRN | P28799 | Progranulin | 1.37 | 0.00 |
| FN3K | Q9H479 | Fructosamine-3-kinase | 1.36 | 0.01 |
| CCN1 | O00622 | CCN family member 1 | 1.34 | 0.00 |
| ECM1 | Q16610 | Extracellular matrix protein 1 | 1.19 | 0.00 |
| PHC2 | Q8IXK0 | Polyhomeotic-like protein 2 | 1.18 | 0.01 |
| CDC42EP2 | O14613 | Cdc42 effector protein 2 | 1.16 | 0.03 |
| ZNF787 | Q6DD87 | Zinc finger protein 787 | 1.14 | 0.01 |
| IRS2 | Q9Y4H2 | Insulin receptor substrate 2 | 1.14 | 0.00 |
| CEBPB | P17676 | CCAAT/enhancer-binding protein beta | 1.10 | 0.00 |
| SERPINE1 | P05121 | Plasminogen activator inhibitor 1 | 1.09 | 0.00 |
| PTX3 | P26022 | Pentraxin-related protein PTX3 | 1.07 | 0.01 |
| FOSB | P53539 | Protein FosB | 1.06 | 0.01 |
| ZFP36L1 | Q07352 | mRNA decay activator protein ZFP36L1 | 1.00 | 0.00 |
| ADM | P35318 | Pro-adrenomedullin | 1.00 | 0.00 |

**Table S4:** Statistical parameters of selected oxylipins and lysolipids which were sig. upregulated upon 2h IL-1ß treatment in comparison to control samples.

| | **2h IL-1ß vs. Con** | | **6h IL-1ß vs. Con** | | **24h IL-1ß vs. Con** | |
|---|---|---|---|---|---|---|
| **ID** | **$log_2$FC** | **adj. p-value** | **$log_2$FC** | **adj. p-value** | **$log_2$FC** | **adj. p-value** |
| PGA3 | 3.25 | 0.00 | 0.08 | 0.88 | -0.07 | 1.00 |
| 15deoxy-PGJ2 | 3.03 | 0.00 | 0.81 | 0.33 | -2.01 | 0.12 |
| LTB4 | 2.35 | 0.00 | 0.42 | 0.12 | -0.07 | 1.00 |
| 5-HEPE | 1.98 | 0.00 | -0.12 | 0.83 | -0.55 | 0.51 |
| 5-HETE | 1.80 | 0.00 | 1.33 | 0.00 | 0.04 | 1.00 |
